# Investigating population continuity and ghost admixture among ancient genomes

**DOI:** 10.1101/2022.12.01.518676

**Authors:** James McKenna, Carolina Bernhardsson, David Waxman, Mattias Jakobsson, Per Sjödin

## Abstract

Ancient DNA (aDNA) can prove a valuable resource when investigating the evolutionary relationships between ancient and modern populations. Performing demographic inference using datasets that include aDNA samples however, requires statistical methods that explicitly account for the differences in drift expected among a temporally distributed sample. Such drift due to temporal structure can be challenging to discriminate from admixture from an unsampled, or “ghost”, population, which can give rise to very similar summary statistics and confound methods commonly used in population genetics. Sequence data from ancient individuals also have unique characteristics, including short fragments, increased sequencing-error rates, and often limited genome-coverage that poses further challenges. Here we present a novel and conceptually simple approach for assessing questions of population continuity among a temporally distributed sample. We note that conditional on heterozygote sites in an individual genome at a particular point in time, the mean proportion of derived variants at those sites in other individuals has different expectations forwards in time and backwards in time. The difference in these processes enables us to construct a statistic that can detect population continuity in a temporal sample of genomes. We show that the statistic is sensitive to historical admixture events from unsampled populations. Simulations are used to evaluate the power of this approach. We investigate a set of ancient genomes from Early Neolithic Scandinavia to assess levels of population continuity to an earlier Mesolithic individual.

## Introduction

Advances in DNA sequencing have led to rapidly increasing numbers of ancient genomes available for demographic inference. Understanding the relationships among such temporally distributed genomes can help reveal historical demographic patterns that would be impossible to detect using modern genomes alone (Sjödin, Skoglund, and Jakobsson, 2014; Malmström et al., 2009; Green et al., 2010; Lazaridis et al., 2016; Raghavan et al., 2014; Haak et al., 2015; Rasmussen et al., 2015; Lazaridis et al., 2014; Slatkin, 2016; Schraiber, 2017). In the field of human evolution in particular, ancient genomes have been key to discriminating between models of population continuity, admixture and replacement that have accompanied the emergence and spread of technological innovations, cultures and languages around the world (Lazaridis et al., 2016; Skoglund et al., 2012; Raghavan et al., 2014; Haak et al., 2015; Rasmussen et al., 2015; Olalde et al., 2018).

The primary challenge when investigating population continuity among ancient genomes is that changes in population allele frequencies over time (due to genetic drift) can lead to patterns of genetic differentiation among a temporally distributed sample that obscure continuity. Similar patterns can arise under models of historical admixture, particularly when the source of that admixture is an unsampled, “ghost” population (Beerli, 2004; Slatkin, 2005; Lawson, Dorp, and Falush, 2018). The confounding effects of such structure have been demonstrated using both model-based methods of inference (Städler et al., 2009; Rosen et al., 2018; Mazet et al., 2016) and more qualitative approaches (Lawson, Dorp, and Falush, 2018; McVean, 2009; François et al., 2019). Repeated findings of such unsampled admixture events in the histories of human populations (Green et al., 2010; Reich et al., 2010; Wall et al., 2019; Durvasula and Sankararaman, 2020) has spurred the development of methods that aim to detect and quantify ghost admixture in the ancestries of modern populations (Skov et al., 2018; Wall et al., 2019; Durvasula and Sankararaman, 2020).

A further challenge is that due to the high levels of DNA fragmentation and contamination present (Allentoft et al., 2012; Skoglund et al., 2014b), ancient genomes are frequently sequenced to low coverage, without the depth necessary to confidently assign diploid genotypes (Dabney and Meyer, 2019). In the absence of the information that resides in patterns of linkage among loci, we are often limited to those inference methods based on genetic distances (Wang, Zollner, and Rosenberg, 2012; Verdu et al., 2014), diversity indices (Skoglund et al., 2014a; Diego-Ortega-Del and Montgomery, 2018), or allele frequency-based summary statistics (Patterson et al., 2012). Although the increasing availability of aDNA sequences has led to some temporally aware population genetic methods that explicitly account for the differences in drift expected among temporally distributed sequences (Sjödin, Skoglund, and Jakobsson, 2014; Yang and Montgomery, 2015; Rasmussen et al., 2015; Racimo, Renaud, and Slatkin, 2016; Schraiber, 2017; Silva et al., 2018; François and Jay, 2020), very often such techniques rely on good diploid calls, and therefore fail to take advantage of the large number of low coverage ancient genomes available. Many commonly used methods do not explicitly account for the genetic drift expected among a temporally distributed sample, leading to contradictory or misleading results (Schraiber, 2017). The popular model-based clustering methods STRUCTURE and ADMIXTURE for instance, do not presently account for different temporal sampling schemes, which can result in distorted patterns of shared ancestry when a sample includes ancient genomes (François et al., 2019). The placement of individuals on Principle Components Analysis (PCA) projections has also been shown to reflect both the temporal and geographic distribution of samples (Skoglund et al., 2014a; Novembre and Stephens, 2008). Figure 1 demonstrates the confounding effect of such temporal structure. Datasets of temporally distributed samples were simulated under alternative demographic models of population continuity and historical admixture from an unsampled population, resulting in similar projections. High levels of branch-specific drift separating an ancient from modern populations can make it difficult to identify genuine cases of population continuity (Slatkin, 2016).

**Figure 1.**
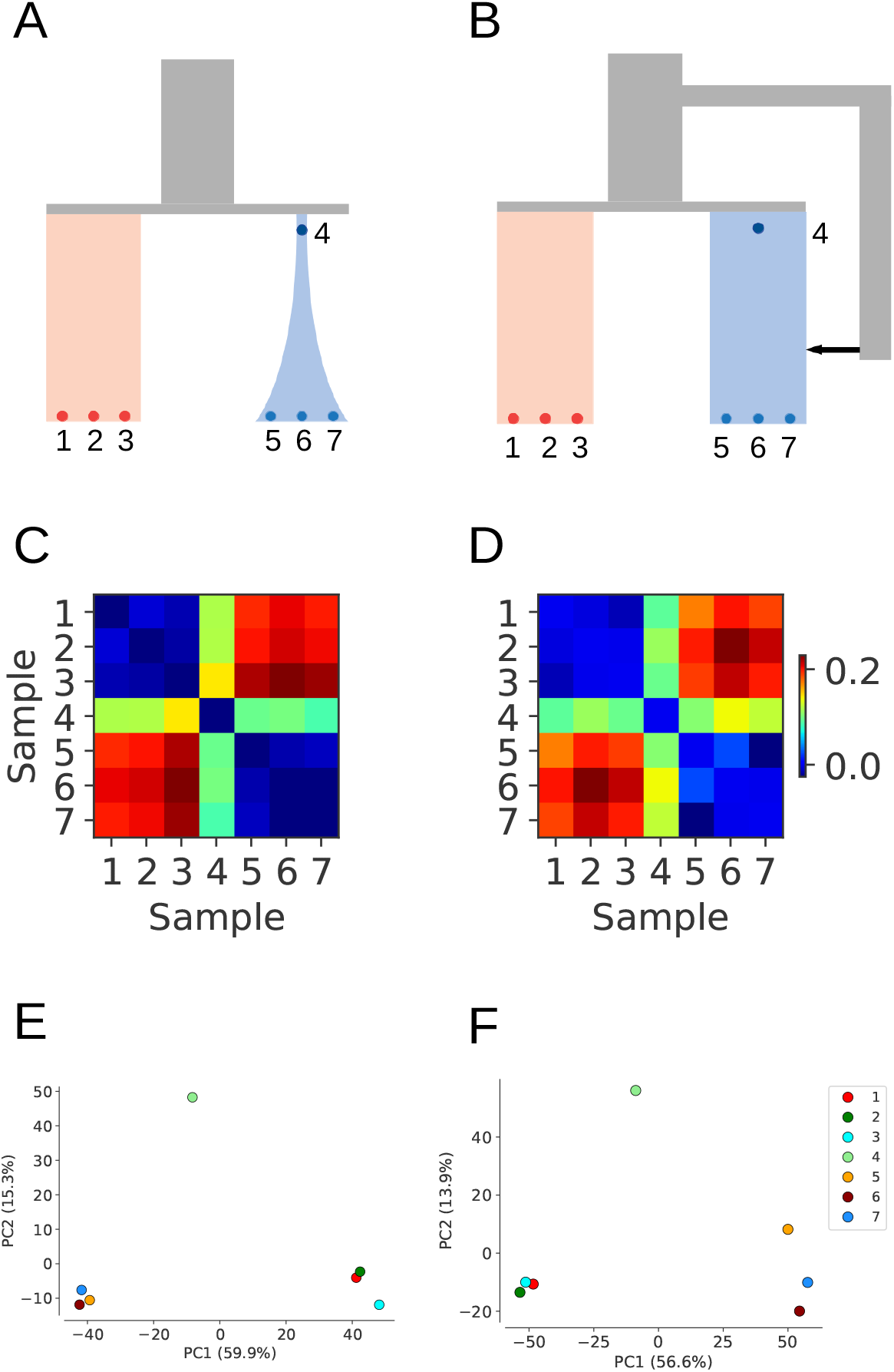
The confounding effects of temporal structure on demographic inference. Datasets simulated under alternative demographic models of A) population branch specific drift, and B) admixture from an unsampled population can result in highly similar pairwise *f*_2_ matrices (C & D) and PCA projections (E & F). In both cases, high levels of drift between ancient and modern samples can obscure relationships of genuine population continuity. Simulations performed using msprime (Kelleher, Etheridge, and McVean, 2016), calculation of *f*_2_-statistics and PCAs performed using scikit-allel (Miles, A. and Murillo, R. and Ralph, P. and Harding, N and Pisupati, R. and Rae, S. and Millar, T., 2010).

Here we propose a novel and conceptually simple approach for investigating population continuity among a temporally distributed sample. The approach is sensitive to historical admixture from unsampled populations. It can further be used with modest coverage ancient genomes and it is robust to missing data. The principle underlying the approach is simple; conditioning on a large number of heterozygote sites in an individual sampled from a population at a particular point in time, the mean proportion of derived alleles at those sites in other individuals is affected differently by the action of genetic drift forwards and backwards in time. The different expectations for these processes allow us to contrast models of population continuity and admixture, and in certain cases, to estimate the proportion of ghost admixture that has occurred. We evaluate the utility and power of this approach using simulations and apply the method to a set of ancient genomes sequenced from two Scandinavian Mesolithic foragers (SHG), five hunter-gatherers from the Neolithic Pitted Ware Culture (PWC) and five contemporaneous farmers from the Funnelbeaker Culture (FBC).

## Method

We will first outline the theory behind the model, followed by the derivation of the central summary-statistic and finally describe a set of simulations that illustrate and evaluate the approach.

### Theory

Although allele frequencies change over time, it is a well known result of population genetics that given a population frequency of *p* of a neutral variant, the expected population frequency of this variant will remain indefinitely at *p* forward in time (Kimura and Ohta, 1969). Consequently, since all (neutral) variants are eventually either lost or fixed in the population, the fixation probability of an allele at present population frequency *p* is *p*. This holds regardless of whether the frequency of the derived or the ancestral variant is considered. On the contrary, the expected frequency of a derived variant looking backwards in time cannot be the same as the present frequency simply because we know that it will eventually disappear (at the time of the mutation that gave rise to it). In fact, we show in the Appendix that the expected frequency of a derived variant *t* generations ago given a present frequency *p* is *pe*^−*τ*^ where *τ* is the scaled time or genetic drift between now and *t* generations ago 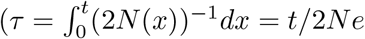, where *N* (*x*) is the diploid population size at time *x* and *N*_*e*_ is the harmonic mean of these *N* (*X*) between *x* = 0,*· · ·, t*). To illustrate, in figure 2, conditional on population frequency *p* in population A, the expected frequency of the derived variant in the branch below population A is *p* (independent of *τ*_1_) while the expected frequency of the derived variant in the right population branch is 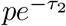 (independent of *τ*_3_).

**Figure 2.**
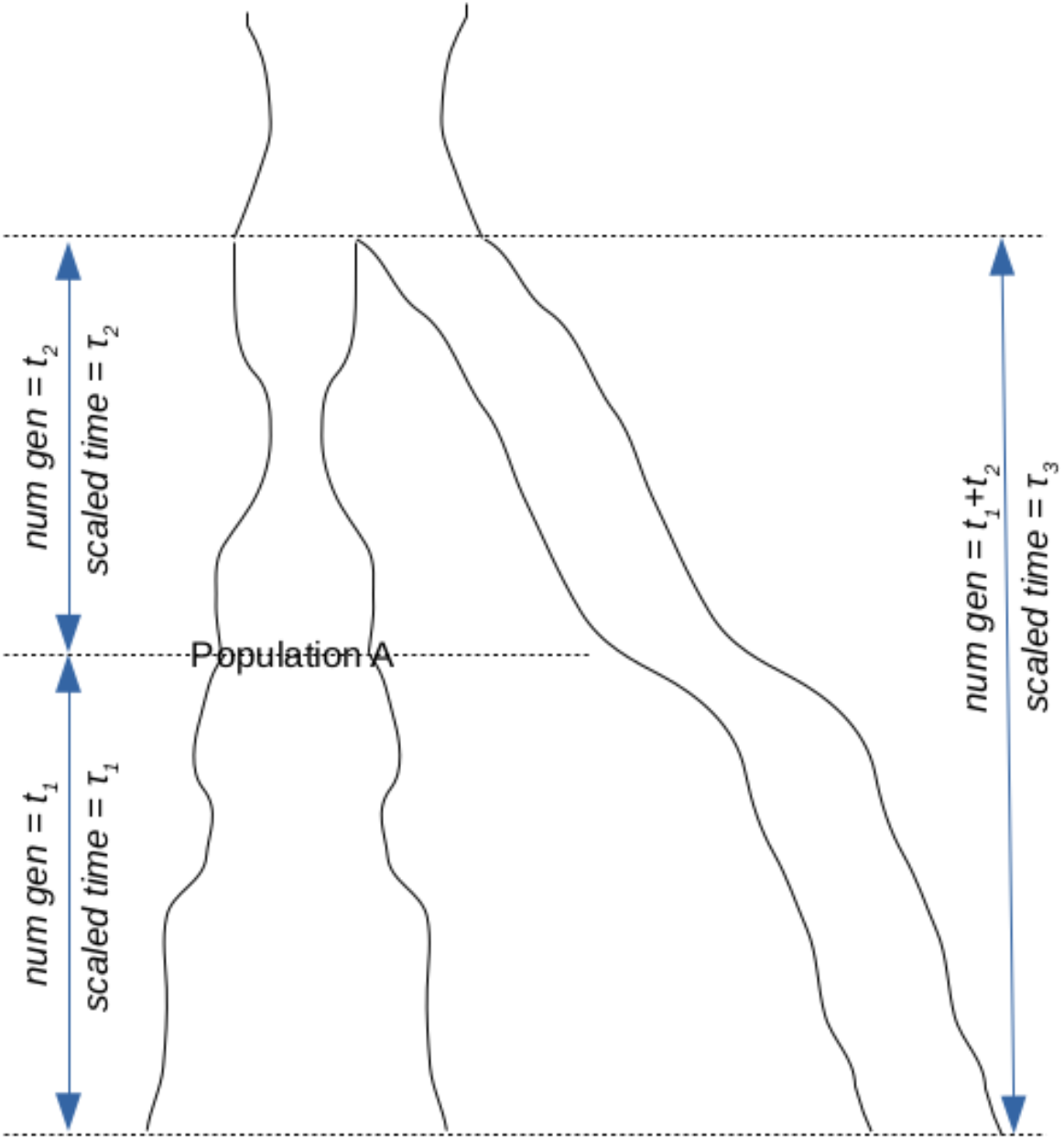
An illustration of the model being analyzed when the two most recent populations are at the same generation. Note however that since scaled time is different from the number of generations, *τ*_3_ may be different from *τ*_1_ + *τ*_2_.

Now consider that an individual (the *anchor individual* for brevity) has been sequenced and genotypes have been called. If we restrict the analysis to heterozygote sites (and we assume that the ancestral and derived variant is known), we know that the derived variant is at least as old as the anchor individual. We also know that it is neither fixed nor lost at this time point which allows us to use the diffusion theory results of the Appendix. The population frequency of the derived variant is not the same at all of these heterozygote sites, but the conditioned distribution has an expected value which we will denote by 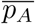 where

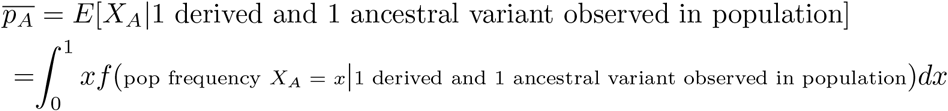

Thus, if we refer to this set of heterozygote sites in the anchor individual as *H*_*A*_, the probability to draw the derived variant at a site in the set *H*_*A*_, in the same population that the anchor individual was sampled from, is 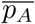. Moreover, for any individual that lived more recent in time and that traces all of its ancestry to population A, the probability to draw the derived variant among *H*_*A*_ is also 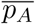. Such an individual can be said to be completely continuous with the population the anchor individual was sampled from.

### The anchor statistic (*R*_*d*_(*A, x*))

Given an anchor individual A and a set *H*_*A*_ of heterozygous sites in A, we define the *anchor statistic* for a test individual *x* as the probability to draw the derived variant in *x* at sites in *H*_*A*_ and refer to this statistic as *R*_*d*_(*A, x*). For empirical data, for low coverage data we do this by adding *k/*(*k* + *l*), where *k* is the number of reads supporting the derived and *l* is the number of reads supporting the ancestral, for all sites with *k* + *l >* 0 and then dividing by the total number of sites (among the *H*_*A*_ sites) with *k* + *l >* 0. For high coverage data we would just use the number of heterozygous sites (*n*_*het*_), the number of homozygous derived sites (*n*_*der*_) and the number of homozygous ancestral sites (*n*_*anc*_) among the *H*_*A*_ anchor sites and do (0.5*n*_*het*_ + *n*_*der*_)*/*(*n*_*het*_ + *n*_*der*_ + *n*_*anc*_).

### Estimating admixture

Consider an individual that traces an amount 1 − *γ* of its genetic ancestry to the anchor population but a proportion *γ* of its genetics ancestry comes from a population that diverged from the anchor population branch *τ* units of genetic drift prior to the anchor population. The average frequency at the time of the split of the latter population (and hence in all of that branch forwards in time) is then 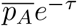 (see above). The probability to draw the derived variant among heterozygote sites in the anchor is therefore no longer 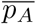 but

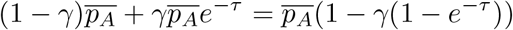

Next, consider the model set-up shown in figure 3(a). If we sample four genomes, (*A*_1_, *A*_2_, *B* and *C*), of which *A*_2_ and *B* are assumed to be more recent but continuous with the population *A*_1_ is sampled from. *C* is assumed to be sampled more recently than the others and the admixture event. The admixture event comprises a proportion *γ* from a population diverging *τ* time units back from *A*_1_, and *τ* + *τ*_*A*_ drift units back from sample *A*_2_.

**Figure 3.**
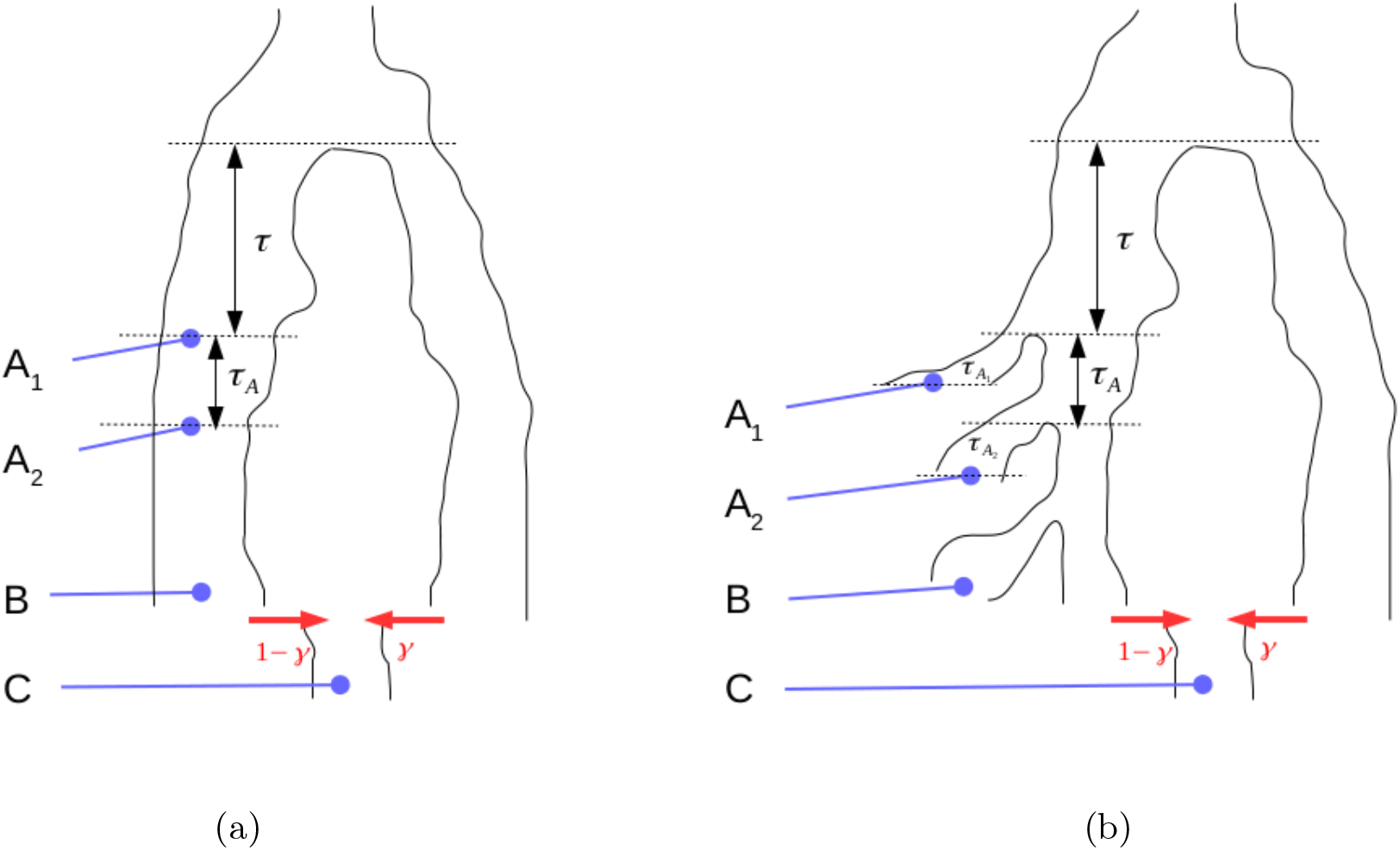
Set up for estimating proportion of admixture (*γ*) from a population diverging *τ* time units before anchor sample *A*_1_.

If by *E*[*R*_*d*_(*A, x*)] we denote the expected value of the anchor statistic for a test individual *x* among sites that are heterozygote in an anchor individual *A*, then

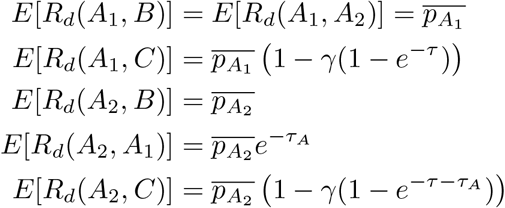

Furthermore, if we allow private drift to each of the sampled individuals so that, in effect, there is no contribution from any of the possible anchor populations to any of the possible test individuals, (see figure 3(b)) as well as sequencing errors specific to the anchors (denoted by 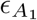 and 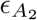) we have

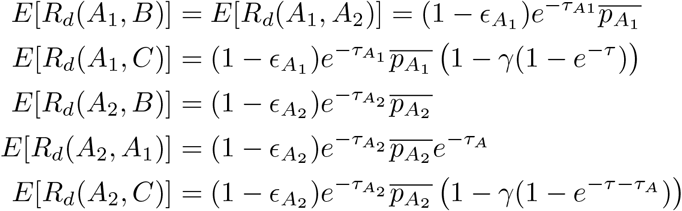

In effect, this is the same as a rescaling of 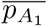 and 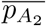 and for both figure 3(a) and figure 3(b) we have

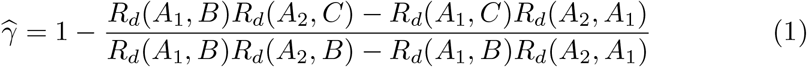

Where 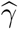 is an estimate of the proportion of ancestry in *C* from the “ghost” population to the right in figure 3.

### Simulations

#### Detecting population continuity and quantifying admixture

Here we use simulations to investigate the power of this approach to detect population discontinuity and to quantify admixture from an unsampled population. Datasets were simulated under alternative demographic models of population continuity (figure 4A), and population discontinuity including two pulses of admixture from a population diverging 4000 generations ago (figure 4B). All simulations were performed using msprime (Kelleher, Etheridge, and McVean, 2016), aggregating data across 1000 runs of a 2M base sequence.

**Figure 4.**
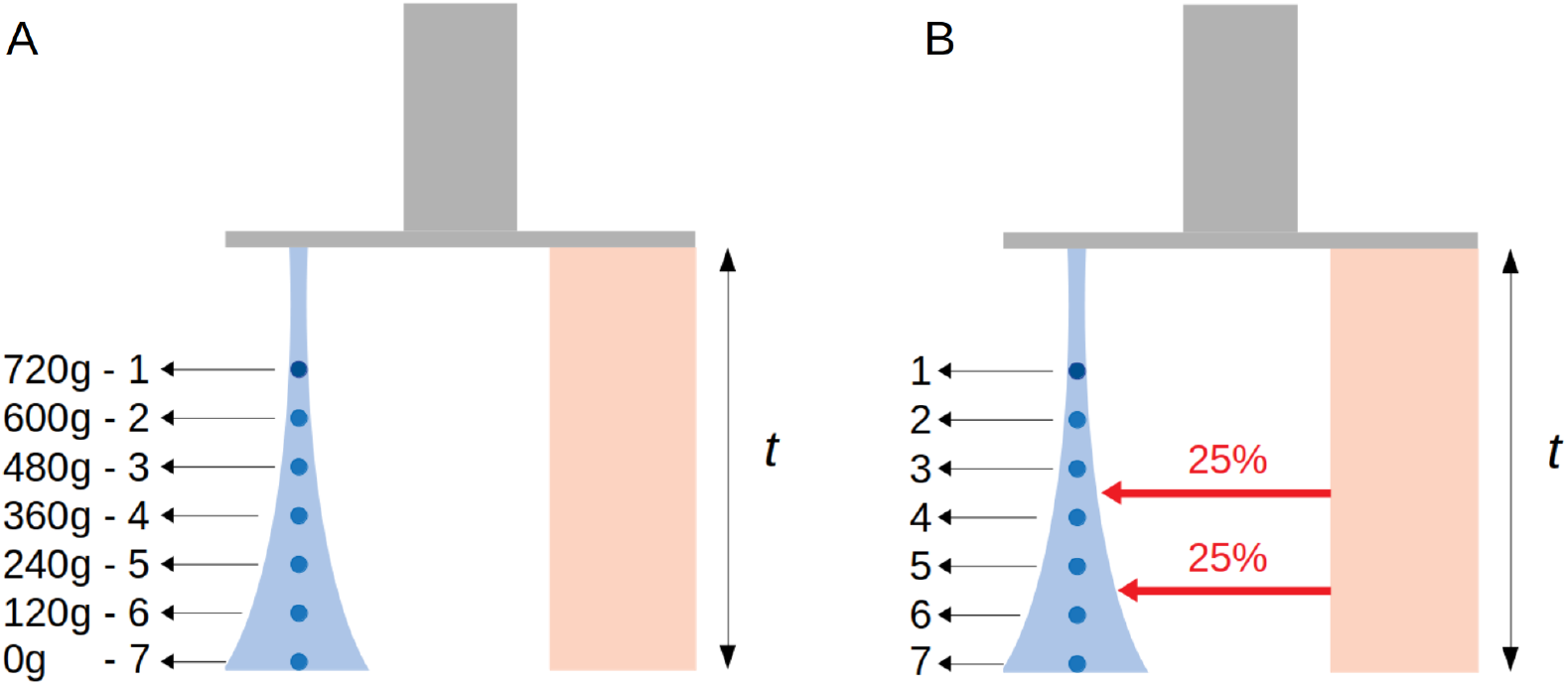
Illustrating the set-up used for simulation of data under models of A) population continuity and B) pulsed admixture from an unsampled population. Samples 1 *−* 7 are taken at different time points from one population undergoing a population expansion from initial size at population divergence of 1, 000 diploid individuals, to final size at present day of 10, 000. Times shown in generations with 1 gen=29 years. Population divergence time *t* = 4, 000 generations.

In each model, 7 diploid individuals were sampled at times ranging from present day to 720 generations in the past. The generation time was set to 29 years, with mutation and recombination rates fixed at 1.45*×* 10^*−*8^ and 1.25 *×*10^*−*8^ respectively. Both models include a population divergence event 4, 000 generations in the past, with a population experiencing an expansion from *N* = 1, 000 at divergence time to *N* = 10, 000 at present day. All other branches of the models are fixed and constant at *N* = 10, 000. Note however that the anchor framework requires no assumption of fixed or constant population sizes through time to detect population continuity or ghost admixture. Model B in Figure 4 includes two pulses of 25% admixture from the diverging population at 180 and 420 generations respectively. We can condition on heterozygote sites in the oldest individual from our sample (individual 1 sampled at 720 generations) in both models, and compare the mean proportion of derived at those sites in more recent individuals.

In order to evaluate the power of this framework to estimate admixture proportions from an unsampled population, a dataset was simulated under the demographic model of a pulse admixture event shown in figure 5. Simulations were performed in an identical manner as before, but this time with four individuals (*A*_1_, *A*_2_, *B* and *C*) sampled at 1, 000, 100, 50 and 0 generations respectively. A single pulse of admixture into this temporal sample was included at 25 generations in the past (between sampling times of individuals *B* and *C*). The performance of in estimating admixture proportions was assessed when three parameters of this model were allowed to vary; admixture proportions (*γ*) varying from 0% to 90%, population divergence time *t* ranging from 1, 000 to 5, 500 generations (genetic drift between 0.05 to 0.2), and the degree of drift in the branch separating anchor samples (number of generations separating *A*_1_ and *A*_2_ varied from 100 to 900 generation which together with a diploid population size of 10, 000 gives that the genetic drift varied from 0.005 to 0.045).

**Figure 5.**
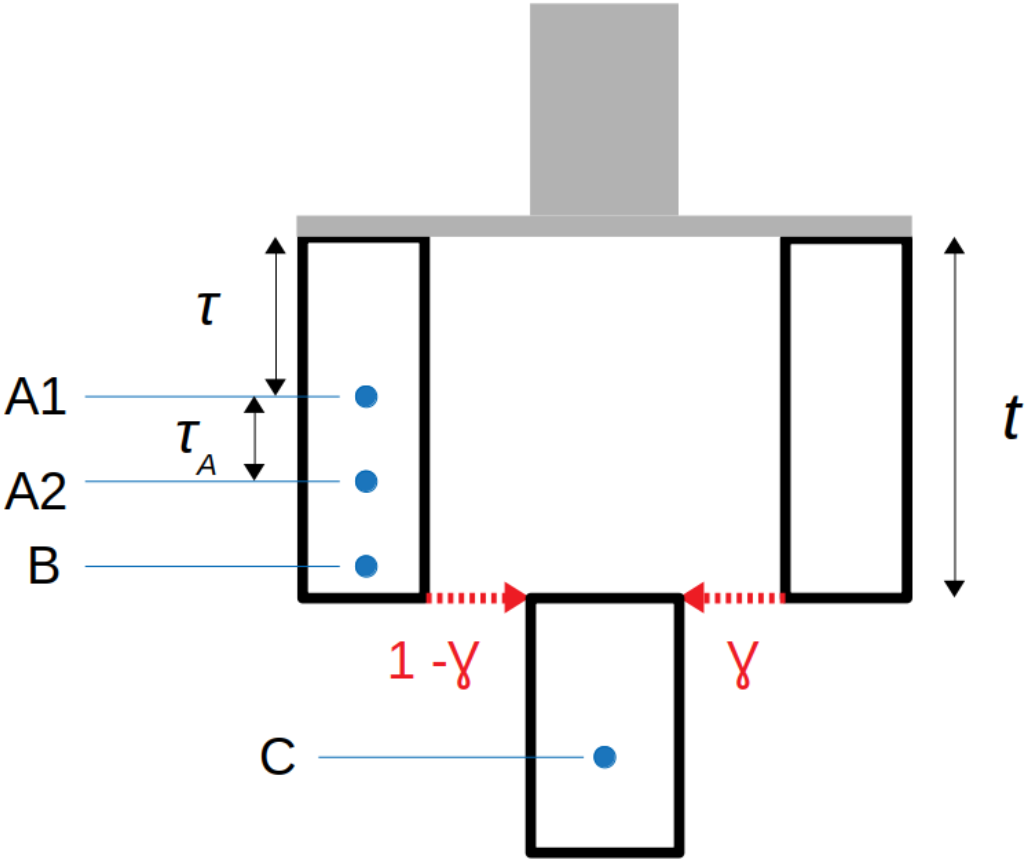
Simulation model set up for estimating proportion of admixture (*γ*) from a population diverging *τ* time units before anchor sample *A*_1_.

## Results

### Simulations

#### Population continuity and ghost admixture

Figure 6 shows the mean proportion derived variants in individuals more recent than the anchor-individual (*R*_*d*_(*Anchor, test*)), for simulated data from models of population continuity and admixture. Forward in time, the mean proportion derived variants remains constant under a model of population continuity even in the presence of strong branch specific genetic drift followed by population expansion (figure 4A). Under a model of historical admixture however (figure 4B), there are two clear successive reductions in the mean proportion derived, each associated with separate admixture events. The magnitude of the reduction in proportion derived decreases with the degree of drift along the branch between the anchor-individual (individual 1) and the population divergence event (*t* = 4, 000) with the branch leading to the population that mixes into the sampled population.

**Figure 6.**
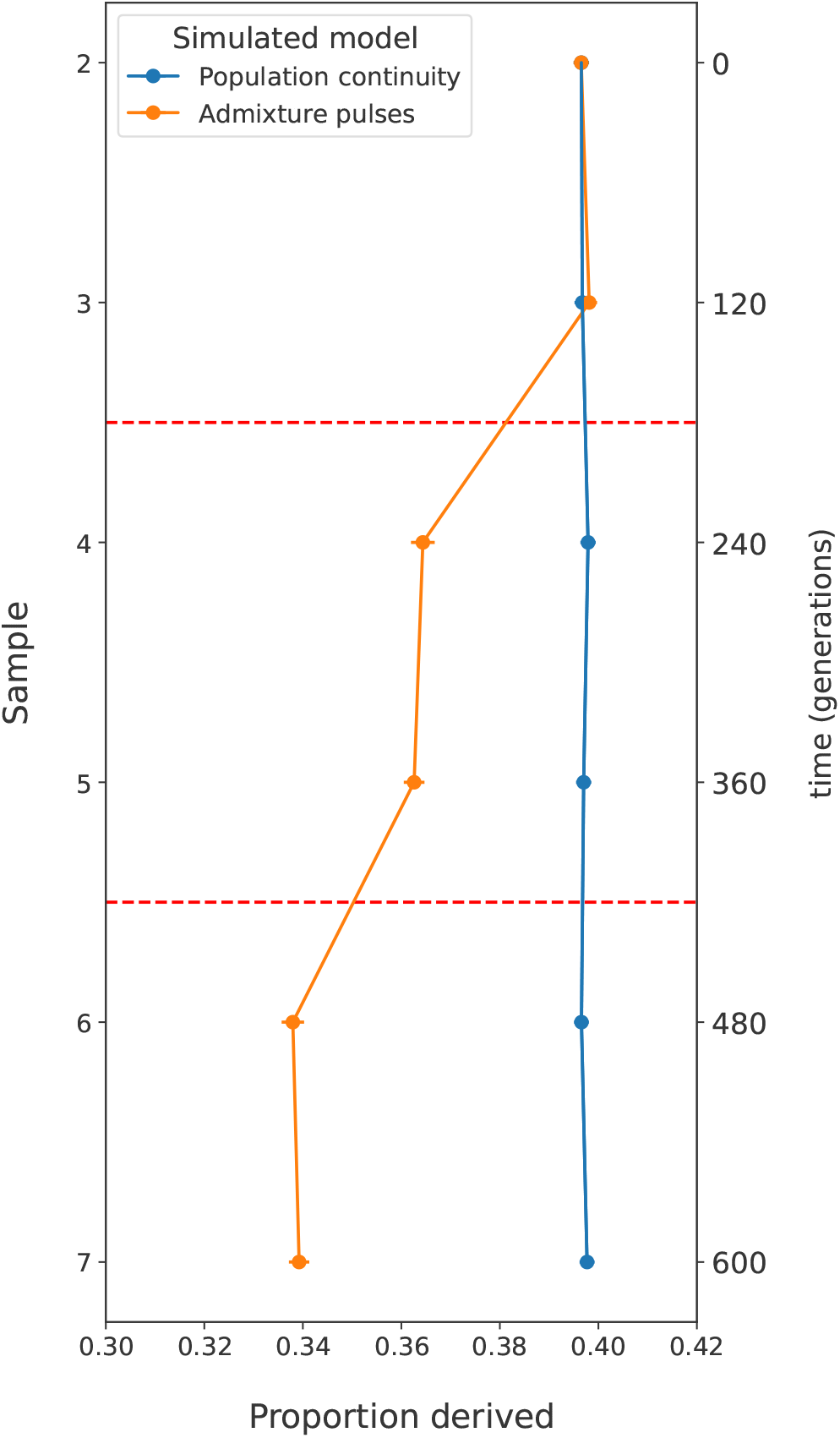
Anchor statistic under simulated models of population continuity and admixture. In each case the oldest sampled individual (1) is used as the anchor-individual, with the mean proportion derived at anchor heterozygote sites counted in all more recent individuals. Red dashed lines at *t* = 180 & *t* = 420 generations indicate admixture events (25%) in the Admixture Pulse model. The mean proportion derived variants remains constant forwards in time for individuals sampled from a model of population continuity. The mean proportion derived variants shows successive reductions for individuals sampled from a model that includes admixture. Error bars show two weighted-block jackknife standard deviations.

#### Estimating admixture proportions

The anchor statistic can also be used to estimate admixture as outlined in the Methods section. From simulations, we show that our estimate of admixture proportion is highly correlated to the true admixture proportion (*r* = 0.98, Pearson correlation). Furthermore, the accuracy of the admixture-estimate does not appear to be affected by the true proportion of admixture itself (figure 7a). Varying the population divergence time (*t*) from 1000 to 5500 generations does not change to accuracy of the estimation (figure 7b). Figure 7c shows that admixture estimation uncertainty increases as the degree of drift between anchor-individuals is reduced from 0.045 to 0.005 in drift units. These observations suggest that it is not the genetic drift prior to the anchor-individuals, but rather the degree of drift between the two anchor samples that influence accuracy of admixture estimation.

**Figure 7.**
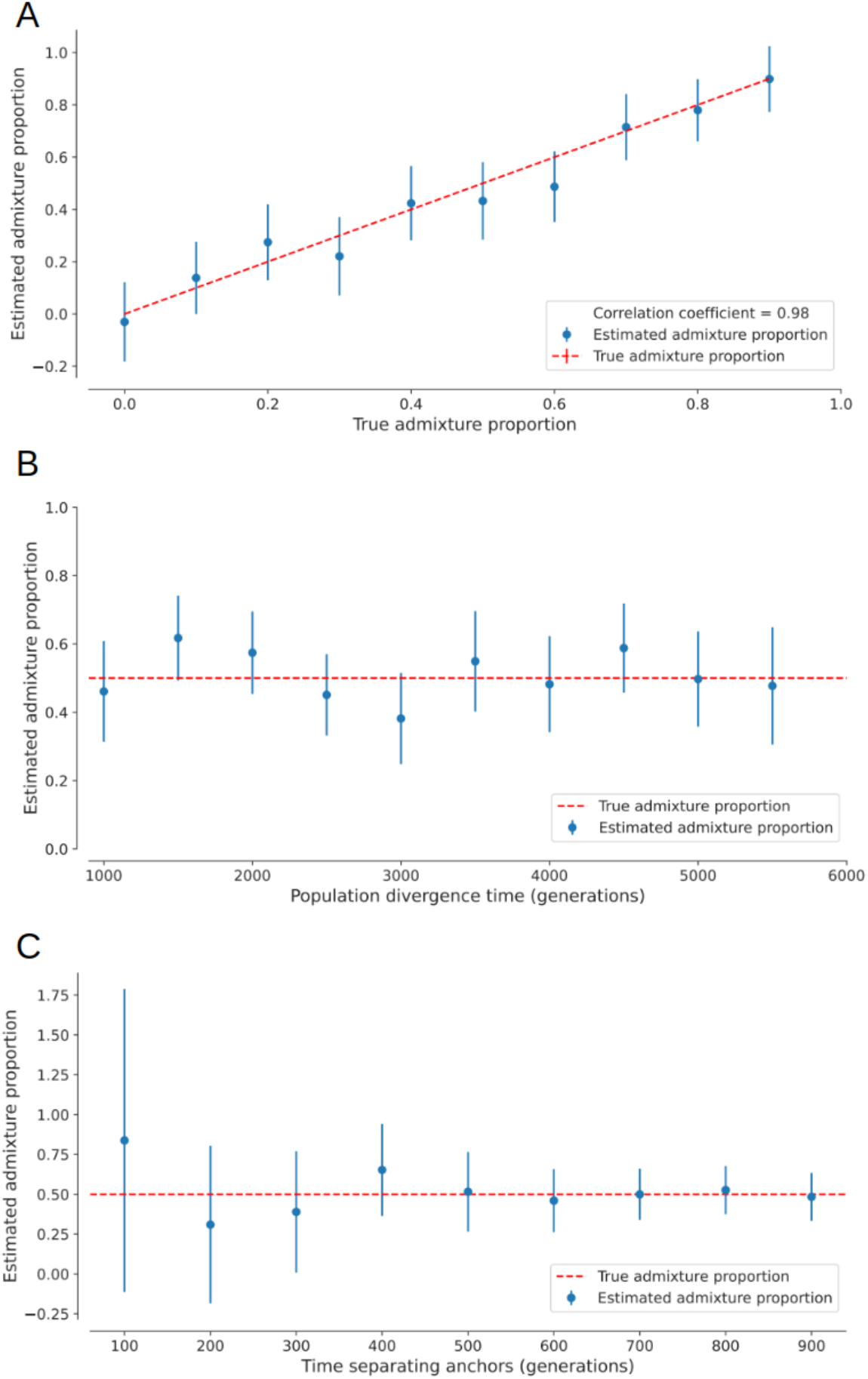
Estimating admixture proportions from an unsampled population with A) a fixed divergence time (*t*) of 4, 000 generations and admixture proportions varying from 0 to 0.9, B) a fixed admixture proportion of 0.5 and *t* varying from 1, 000 to 5, 500 generations, and C) a fixed admixture proportion of 0.5 and *t* of 4, 000 generations, with time separating anchors *A*_1_ and *A*_2_ ranging from 100 to 900 generations. In each case, sampling time of anchor-individual *A*_1_ is 1, 000 generations and population sizes are constant at 10, 000 diploid individuals. Error bars show two weighted-block jackknife standard deviations.

Estimating admixture proportion among a temporal sample therefore requires two individuals of sufficiently high coverage that heterozygotes can be confidently called, and for those anchor individuals to be separated by enough genetic drift to gain the necessary power for accurate estimation.

#### Empirical data

##### *β*-drift and a convenient transformation of *R*_*d*_(*A, x*)

Empirical data will often have a proportion erroneously called heterozygote anchor sites and it is hard to completely rule out some level of private drift associated with the anchor population. For these reasons we argue that studying *−ln*(*R*_*d*_(*A, x*)) is preferable to *R*_*d*_(*A, x*) (where *A* is an anchor and *x* is any test individual. The incentive for this is that the distance between different estimates then has a clear biological interpretation as the difference in backward drift in the path from the anchor population to the population of a test individual. For brevity, we will refer to this drift as *β*-drift which is always relative to some anchor population. To illustrate, consider two test individuals *B* and *C* with *β*-drift *τ*_*B*_ and *τ*_*C*_ relative to anchor *A*. If *ϵ*_*A*_ represents the probability of an anchor site being due to an error in the anchor sequence and not representing a a true heterozygote site and *τ*_*A*_ the amount of private drift associated with the anchor, then

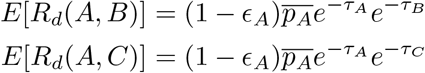

so that

*−ln*(*R*_*d*_(*A, C*)) + *ln*(*R*_*d*_(*A, B*)) is an estimate of the difference in *β*-drift between *C* and *B*: *τ*_*C*_ − *τ*_*B*_

##### Investigating population continuity from Mesolithic foragers to different Neolithic groups in Scandinavia

Scandinavia as a region holds special interest when considering the roles that population continuity and admixture have had on the emergence and spread of human cultures. It was the last region of Europe to become free of ice after the Last Glacial Maximum, and it harboured some of the last populations of hunter-gatherers in Europe. The earliest expressions of Neolithic material culture also emerged relatively late, with the arrival of the Funnelbeaker culture (FBC) circa 3500BCE to southern Scandinavia (Midgley, 1992; Lillie et al., 2000; Skoglund et al., 2012; Fraser et al., 2018). Paleogenetic data show a close connection between these early farmers of Scandinavia and other Early and Middle Neolithic farmers of Europe (Malmström et al., 2015; Malmström et al., 2019), providing strong support for a model of population discontinuity between the Scandinavian hunter-gatherers of the Mesolithic and the arriving farmers. The Neolithic Pitted Ware Culture (PWC) represents an intriguing case of a hunter-gathering culture in Scandinavia that emerge circa 3200BCE and post-dates the migration of the farmers that spread the FBC culture into Scandinavia. Despite this, the results of several studies have indicated that the PWC culture predominantly shows genetic affinity to earlier Mesolithic hunter-gatherers rather than contemporary Neolithic farmers (Skoglund et al., 2012; Skoglund et al., 2014a). Furthermore, archaeological data has been used to paint a picture of two cultures overlapping in both time and space, coexisting in parallel for several hundred years and yet still maintaining distinct material cultures and dietary patterns, in addition to maintaining distinct genetic make-ups (Malmström et al., 2009; Fraser et al., 2018; Coutinho et al., 2020; Malmström et al., 2019).

We assembled a panel of fully UDG (uracil-DNA glycosylase) treated high-coverage ancient genomes from an ongoing effort to increase genome coverage from Scandinavian Stone Age humans, including a Mesolithic hunter-gatherer from the Baltic Island Stora Karlsö (sf12) and five PWC hunter-gatherers from the Baltic island of Gotland, together with five FBC farmers from the Megalithic passage tombs of Gökhem and Rössberga. In order to evaluate the impact genome coverage has on the statistics, we also included two relatively low coverage blunt-end screening genomes. One of these comes from the same ancient remains as one of the high-coverage PWC samples (ajv058) while the other one (sf9) is contemporaneous with and from the same site as sf12 (supplementary table 1, Günther et al., 2018; Fraser et al., 2018; Coutinho et al., 2020; Malmström et al., 2019). Although from distinct material cultures, all PWC and FBC individuals are broadly contemporaneous from the same period of the Nordic Middle Neolithic (circa 3200 *−* 2300BCE). The raw sequence data was processed according to Alves et al. (2022) and Günther et al. (2018). See Appendix section “Data processing” for how genotypes were called.

**Table 1:**
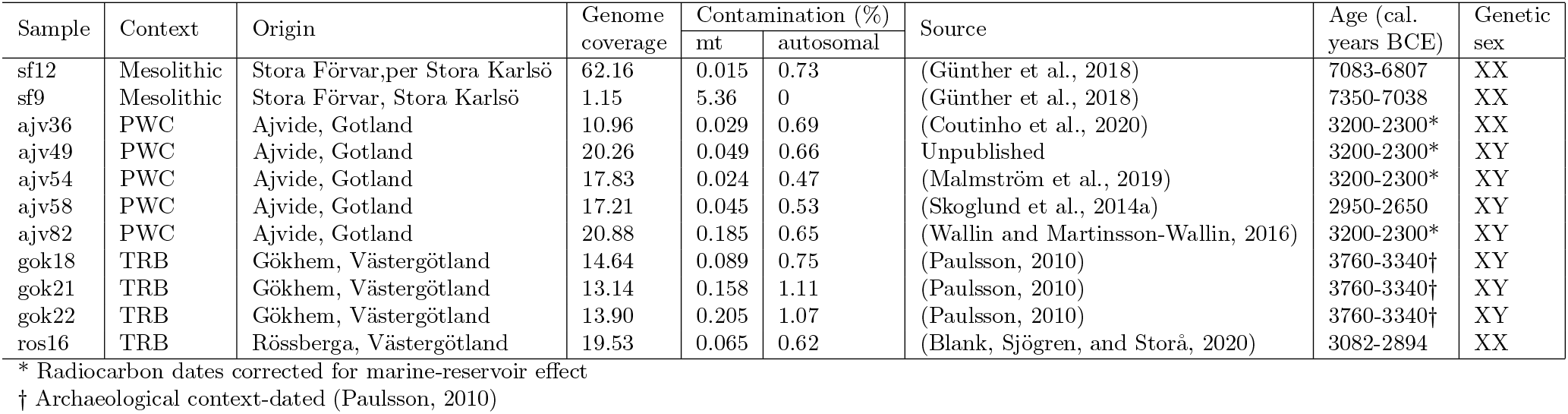
Information on the 10 individuals used in this study. mt, mitochondrial; cal, calibrated; BCE, before common era; PWC, Pitted Ware Culture; TRB, Funnel Beaker Culture.

The Mesolithic hunter-gatherer sf12 was used as the anchor-individual so as to measure differences in *β*-drift between this individual and a set of other, more recent, individuals (figure 8B). All PWC individuals show a similar mean *β*-drift that is lower compared to the FBC individuals but higher than for sf9. These results are consistent with previous findings of modest gene-flow between the contemporaneous PWC and FBC groups (Malmström et al., 2009; Fraser et al., 2018; Coutinho et al., 2020) but also that PWC is not a direct descendent population from SHG.

**Figure 8.**
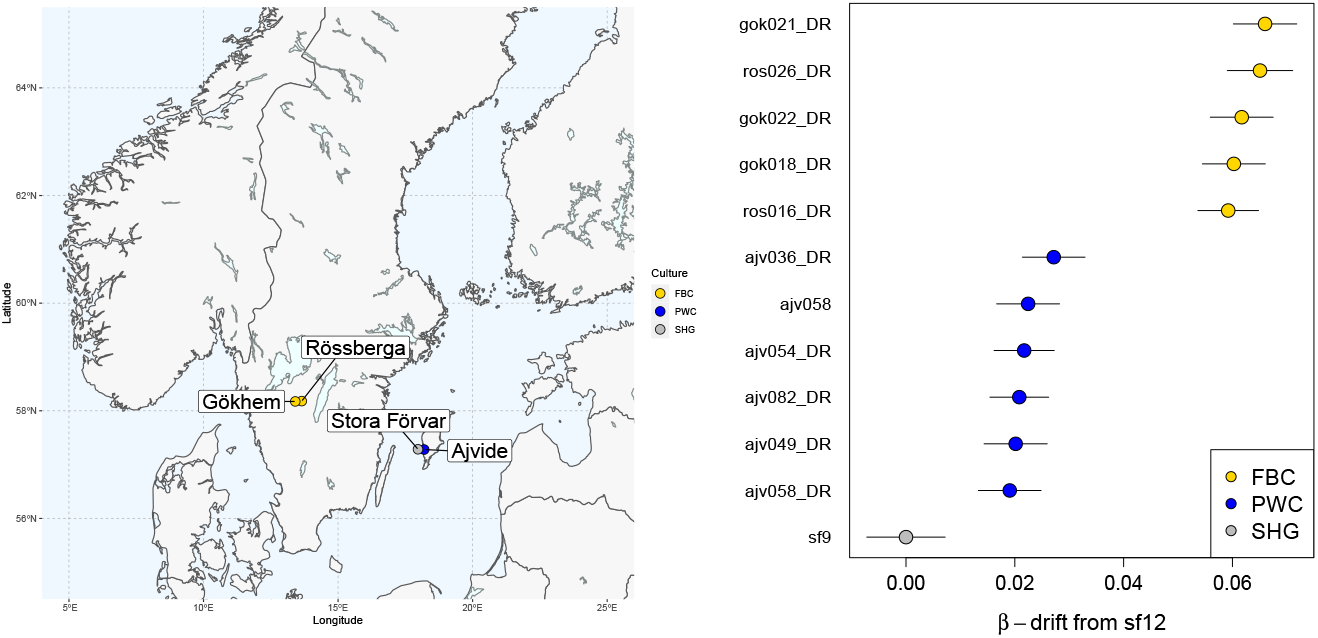
A) Map showing distributions of the sampled individuals that were either representatives of the Funnel Beaker culture (FBC) or the sub-Neolithic Pitted Ware culture (PWC) during the Nordic Middle Neolithic period. All PWC individuals from Ajvide burial, Gotland. FBC burial sites sampled include Gökhem passage grave, and Rössberga passage grave. Mesolithic individuals sf9 and sf12 from Stora Förvar on Stora Karlsö. B) *β*-drift relative to sf9 for different individuals with Scandinavian Mesolithic hunter-gatherer sf12 used as anchor. Error bars show two weighted-block jackknife standard deviations. The suffix “DR” in the sample names indicate damage-repair through full UDG treatment of the sequenced libraries.

## Discussion

Ancient genomes have the capacity to revolutionize our understanding of the demographic processes contributing to patterns of genetic variation among current-day humans as well as among other species. aDNA can reveal population continuity through time and aid in the detection of historical admixture events and population replacements. Particularly in studies of human demographic history, aDNA has proven an important resource in understanding the pre-historic movements of people that have spread cultures, languages, and technologies to new areas (*e*.*g*. Arcos and Raghavan, 2023). In order to take full advantage of this resource however, it is essential to develop population genetic tools capable of utilizing samples of temporally distributed genomic data. Here we have outlined a novel and conceptually simple approach that is capable of elucidating questions of population continuity through time. It is sensitive to admixture from unsampled (“ghost”) populations and can take advantage of the increasing numbers of low coverage genomes available.

Although the concept of continuity is generally clear from the context (*e*.*g*. Mattila et al., 2023), for the purpose of discussing the relationship between the anchor statistic and continuity, we will with a statement such as “an individual *x* has continuity level *c* with population A (that existed at time point *t*)” mean that a proportion *c* of all ancestors to *x* living at time point *t* belonged to population A. This in turn implies that a proportion *c* of *x*’s autosomal genetic material is expected to trace back to individuals living in population *A*. With respect to the different scenarios in figure 9, this definition would imply that all test individuals in scenarios a) and b) have continuity level 1 with the anchor population while scenario c) has continuity level 0. In d), all test individuals would have some continuity level *c* while only the two most recent test individuals in scenario e) would have a level of continuity larger than 0. In contrast, all test individuals in each of the scenarios *could* have identical amounts of *β*-drift relative to the anchor. Hence, unless *β*-drift is 0, it can be difficult to assess any level of continuity at all based exclusively on the anchor statistic. However, when it can be assumed that one test individual has no *β*-drift (*i*.*e*. it has continuity level 1 with the anchor population) the absolute *β*-drift can be assessed. Since *sf* 9 was sampled from the same location as sf12 and both were estimated to be around 7, 000 years old (see table 1 in the Appendix), we assumed that sf12 and sf9 were drawn from the same population and could therefore translate values of the anchor statistic to actual *β*-drift from the population sf12 and sf9 were sampled from. From this analysis we conclude that although the PWC individuals have about 0.02 of *β*-drift relative to SHG (represented by sf12 and sf9 in this analysis) and thus that they have ancestry in other populations than SHG that were contemporaneous with SHG. Any admixture event with proportion *c* with a population that has an additional *τ* amount of *β*-drift such that −*ln*(*ce*^−*τ*^ + 1 − *c*)*≈* 0.02 would give this pattern. This is consistent with that they have none of their ancestry tracing back to SHG but to a population that diverged from SHG approximately 0.02 units of drift before the sampled individuals in this study. Alternatively, they may have a significant part of their ancestry in SHG but that they are admixed with populations that diverged considerably more than 0.02 units of drift before the population represented by sf12 and sf9. What is clear is that the FBC individuals have an additional 0.04 amount of *β*-drift from SHG compared to the PWC individuals.

**Figure 9.**
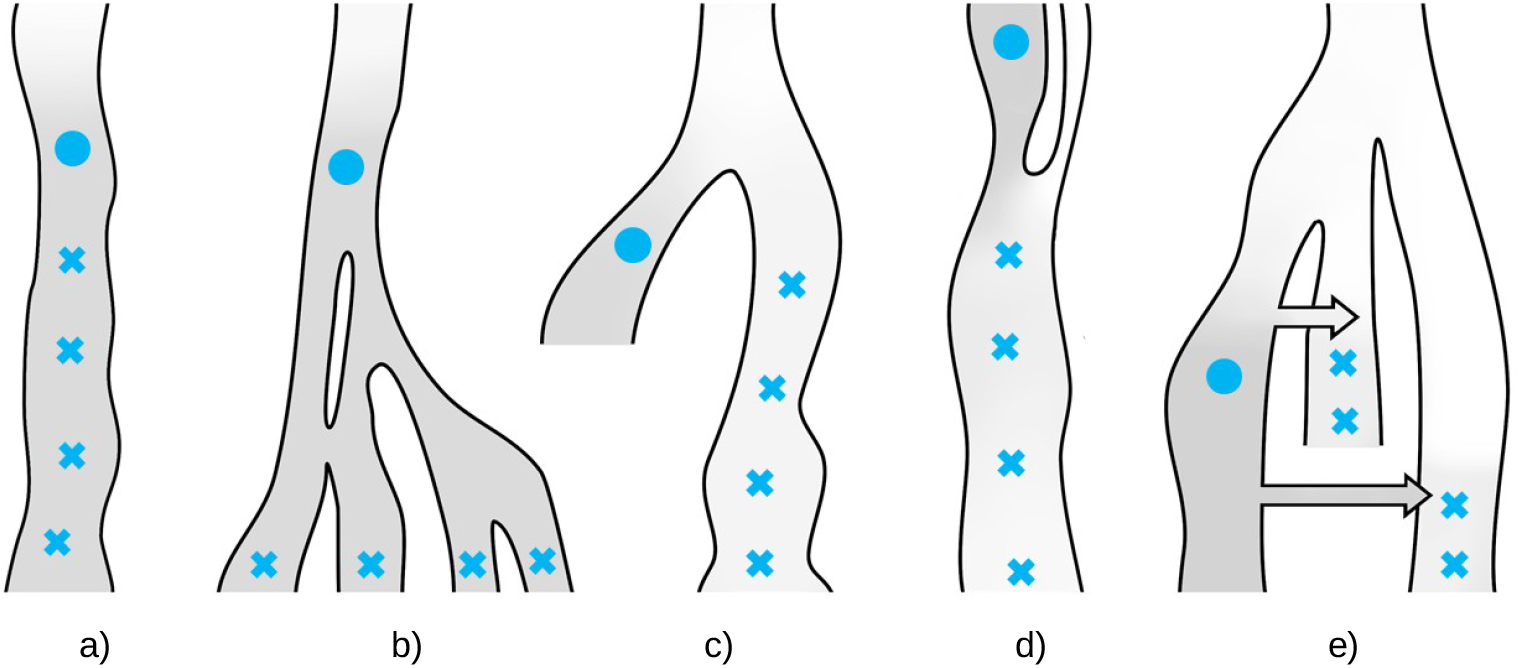
In each of the scenarios a)-d) all test individuals (blue crosses) have the same expected value of the anchor statistic when calculated relative to the anchor (blue filled circle). e): although it will depend on the details of the two admixture events (the arrows), this scenario can result in all test individuals (blue crosses) having the same expected value of the anchor statistic. In all scenarios, the darker the grey, the lower the backward drift in the average path from the anchor population.

We have shown, through simulations and an empirical example, that this approach has the power to infer population continuity and estimate proportions of ghost admixture. Although we did not find a suitable empirical example to estimate proportions of ghost admixture it is a UNIQUE(?!) and potentially powerful aspect of our approach as the method does not require any modeling of the source of the admixture by substitute populations.

Finally, although our set-up is closely related to an outgroup-f3 if the outgroup is chosen to be another species (the same species that is used to call ancestral and derived state for the anchor analysis), the two statistics are not identical and our statistic has the advantage of being an estimate of a biologically meaningful parameter (*β*-drift). As such, our setup is in our view a more natural way of investigating continuity and relationships among ancient samples.

## Acknowledgement

We want to thank Olaf Thalmann, Karl-Heinz Herzig, and Jarosolaw Walkowiak for access to unpublished data, Nikola Vukovic for help with graphic design and Imke Lankheet for Latex assistance. This work was supported by the Knut and Alice Wallenberg foundation. The computations and data handling were enabled by resources (SNIC2022/2-11 and p2018003) provided by the Swedish National Infrastructure for Computing (SNIC) at UPPMAX.

## Appendix

### Moments for the population frequency

For brevity, we suppress the dependence of *p* and *τ* in the notation and write 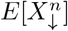 for the *n*’th moment of the population frequency after *τ* units of genetic drift *forwards* in time conditional on population frequency *p* and 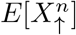 for the *n*’th moment of the population frequency going back *τ* units of genetic drift conditional on population frequency *p*. These moments are derived below.

### Moments for *X*_*↓*_

To derive the moments of *X*_*↓*_, note that given a population frequency *x* of the derived variant, the probability of obtaining a sample of size *n* with only the derived allele is *x*^*n*^. Thus, 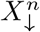 is the probability to pick only the derived variant in a sample of size *n*, and 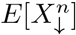 is the expected value of this probability. Conditional on the frequency of the derived variant being *p, τ* time units ago, the expected probability for such a sample is derived by averaging over the number of ancestors to the sample *τ* time units ago, where each of those ancestors need to be of the derived type. Hence, 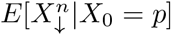, or the expected probability to observe only the derived variant in a sample of size *n* is

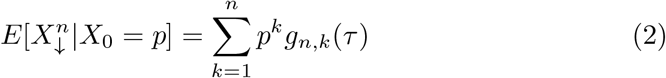

with *g*_*n,k*_(*τ*) being the probability of there being *k* ancestors at time *τ* to a sample of *n* gene-copies (a much longer recursive proof using diffusion theory can be obtained from the authors upon request). Specifically

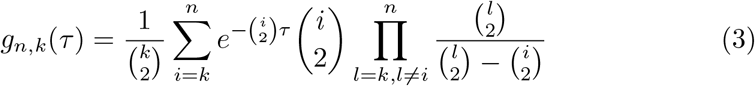

for 2 *≤ k < n* with special cases 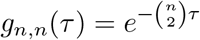 and 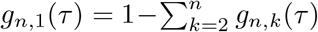 (Tavaré, 1984; Wakeley, 2009).

Since *g*_1,1_(*τ*) = 1, *g*_2,2_(*τ*) = *e*^*−τ*^, *g*_2,1_(*τ*) = 1 *− e*^*−τ*^ we have

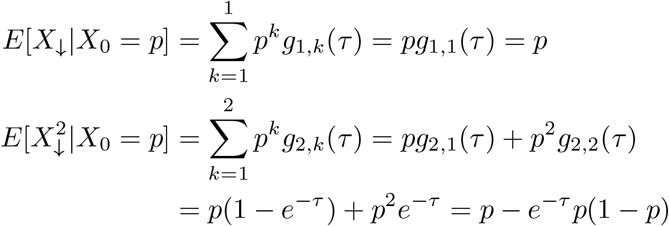

### Moments for *X*_*↑*_

Define *ϕ*(*x, τ, p*) to be the probability density of frequency *x* at time *τ*, given that *X*_*A*_ = *p*. The density backward in time is, in fact, the same as the forward density conditional on extinction (since by definition the derived variant goes extinct backwards in time) (Griffiths, 2003). The forward density conditional on the allele ultimately going extinct is (using *\** to denote conditional on extinction)

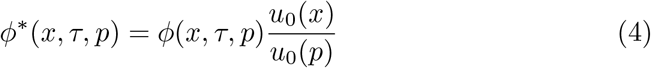

where *u*_0_(*z*) is the probability of an allele at frequency *z* ultimately being lost. In a neutral model, *u*_0_(*z*) = 1 *− z*. We get

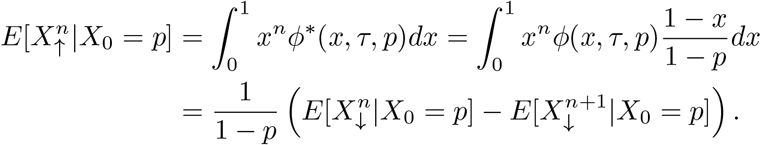

For *n* = 1 we have

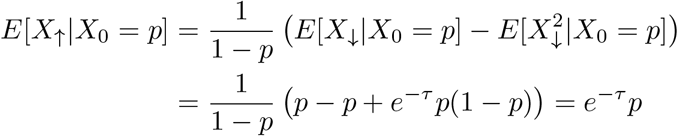

### Sample metadata

#### Data processing

Before calling, a base quality score recalibration (BQSR) step was performed on the 5 terminal bases of each read by reducing the quality of all 5’ T:s and all 3’ A:s to phred-score 2 (*♯*), using a custom python script. Dummy read groups (RG) were added with Picard v 1.118 (ref), followed by indel realignment using known indel sites in the 1000g phase1 data set (ref) with GATK v. 3.5.0 (ref).

A site in the anchor was chosen as an anchor position if 1) the read coverage was neither within the 5% lower or 5% upper tail of the coverage distribution 2) two variants observed at the site and the minor allele frequency was at least 1*/*3, 3) one of the variants could be confidently called as the ancestral variant (in our case, we demanded that all three apes had the same variant (without missingness) and that this variant was one of the variants observed in the anchor). Allele counts at anchor positions were parsed using samtools mpileup 1.17 with parameter flags -Q 30 -q 30 (Li, 2011). For site *i* among the anchor positions, the probability to pick the derived variant and the ancestral variant –*p*_*d*_(*i*) and *p*_*a*_(*i*) – for a test individual *x* was calculated. The anchor statistic was then calculate as

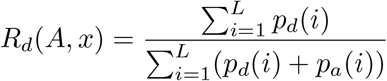

where *L* is the number of anchor positions.

#### Anchor statistic and *f*-statistics

All scripts used for generation of simulated datasets and application of anchor test of population continuity from empirical datasets are available at XXX. Calculation of anchor statistics were performed using custom python scripts and functions. The python program scikit-allel (Miles, A. and Murillo, R. and Ralph, P. and Harding, N and Pisupati, R. and Rae, S. and Millar, T., 2010) was used to calculate *f*2-statistics.

